# When development is constantly but weakly perturbed - Canalization by microRNAs

**DOI:** 10.1101/2021.09.04.458966

**Authors:** GuangAn Lu, Yixin Zhao, Qingjian Chen, Pei Lin, Tian Tang, Zhixiong Tang, Zhongqi Liufu, Chung-I Wu

**Author notes:** These authors contributed equally (GuangAn Lu, Yixin Zhao, Qingjian Chen).

## Abstract

Recent studies have increasingly pointed to microRNAs (miRNAs) as the agent of GRN (gene regulatory network) stabilization as well as developmental canalization against constant but small environmental perturbations. Since the complete removal of miRNAs is lethal, we construct a Dicer-1 knockdown line (*dcr*-*1* KD) in Drosophila that modestly reduces all miRNAs. We hypothesize that flies with modest miRNA reductions will gradually deviate from the developmental norm, resulting in late-stage failures such as shortened longevity. In the optimal culture condition, the survival to adulthood is indeed normal in the *dcr*-*1* KD line but, importantly, adult longevity is reduced by ∼ 90%. When flies are stressed by high temperature, *dcr*-*1* KD induces lethality earlier in late pupation and, as the perturbations are shifted earlier, the affected stages are shifted correspondingly. We further show that the developmental failure is associated with GRN aberration in the larval stages even before phenotypic aberrations become observable. Hence, in late stages of development with deviations piling up, GRN would be increasingly in need of stabilization. In conclusion, miRNAs appear to be the genome’s solution to weak but constant environmental perturbations.

## Introduction

Living organisms face internal and external fluctuations constantly. These fluctuations would broadly perturb many phenotypes. Hence, the maintenance of stable phenotypic output and normal development under such perturbations is an important topic as emphasized by C.H.Waddington (Waddington 1942; Waddington 1959). He coined the term ‘developmental canalization’ for the development along a defined path. A standard depiction is a ball traveling along a canal in a landscape of many canals, each for a tissue of a particular species (Waddington 1957; Scharloo 1991).

There are two approaches to observing developmental canalization. The first one is to measure the variance of phenotypic values at a particular stage of development. Such measurements of phenotypic robustness have been frequently adopted in the literature (Levy and Siegal 2008; Kasper et al. 2017; Hintze et al. 2021; for reviews, please see Rendel 1967; Félix and Barkoulas 2015; Takahashi 2019). A second approach, taken in this study, is to follow the development to the final stages (i.e., maturation to adulthood and death) as Waddington envisaged (Wagner 2005). De-canalization will be observed as increasingly aberrant phenotypes as development progresses.

Molecular biology and mathematical theories of developmental canalization have been proposed (Smith et al. 1985; Wagner 2005; Siegal and Leu 2014; Godin et al. 2020; Saiz et al. 2020; Mirth et al. 2021). While the concept of canalization is developed to account for the phenotypic stability, the effect can be more easily studied at the level of the gene regulatory network (GRN)(Kitano 2004; Guo and Amir 2021). After all, phenotypic changes are downstream of GRN variations. At the molecular level, several genes (Hallgrimsson et al. 2019; Sigalova et al. 2020; Sun and Zhang 2020), e.g., Hsp90 (Rutherford and Lindquist 1998), have been suggested to have the canalizing capacity.

Most notable, a general class of genes that may play a crucial role in canalization has been proposed to be the microRNAs (Hornstein and Shomron 2006; Posadas and Carthew 2014; Liufu et al. 2017).According to the annotations in miRBase (Kozomara et al. 2019), each metazoan species often has several hundred miRNAs and more than 100 of them are expressed in most cells (Ludwig et al. 2016; Zhao et al. 2018; Rahmanian et al. 2019). Each miRNA of ∼ 21 nt in size could weakly but broadly repress hundreds of target genes post–transcription (Bartel 2018).The diffuse (i.e., weak but broad) and uni-directional (downward) regulation is the unique feature of miRNAs (Farh et al. 2005). This mode of regulation has led to the demonstration of incoherent regulation (Liufu et al. 2017) and the mathematical formulation of GRN canalization by miRNAs (Zhao et al. 2017; Chen et al. 2019).

In testing the hypothesis of miRNA canalization, miRNAs collectively are regarded as a single regulatory unit. We then reduce the entire miRNA pool modestly and evenly by creating a *dcr-1* KD line (for Dicer knock-down). In this *dcr-1* KD background, several expectations can be formulated. First, fly development would deviate gradually from the set course as the development progresses. Second, the deviation would be hastened by environmental stresses including temperature aberrations. Third, at the molecular level, *dcr-1* KD should have only marginal effects on transcriptome under normal condition while, under stressful conditions, *dcr-1* KD would greatly exacerbate the effects of the stress. Our study will test if these expectations are fulfilled.

## Results

### PART I –Reduction of total miRNAs by *dcr-1* KD

Here, we reduced the amount of all miRNAs by knocking down the Dicer-1 gene (*dcr-1* KD). We used a weakly and broadly expressed Gal4 construct (Hrdlicka et al. 2002) to drive the interfering RNA that knocks down *dcr-1*(Ni et al. 2011). In RT-qPCR, the observed *dcr-1* expression decreases modestly by 25-30% (Student’s *t* test, *p*<0.05) (**Fig.1A**).

**Figure 1.**
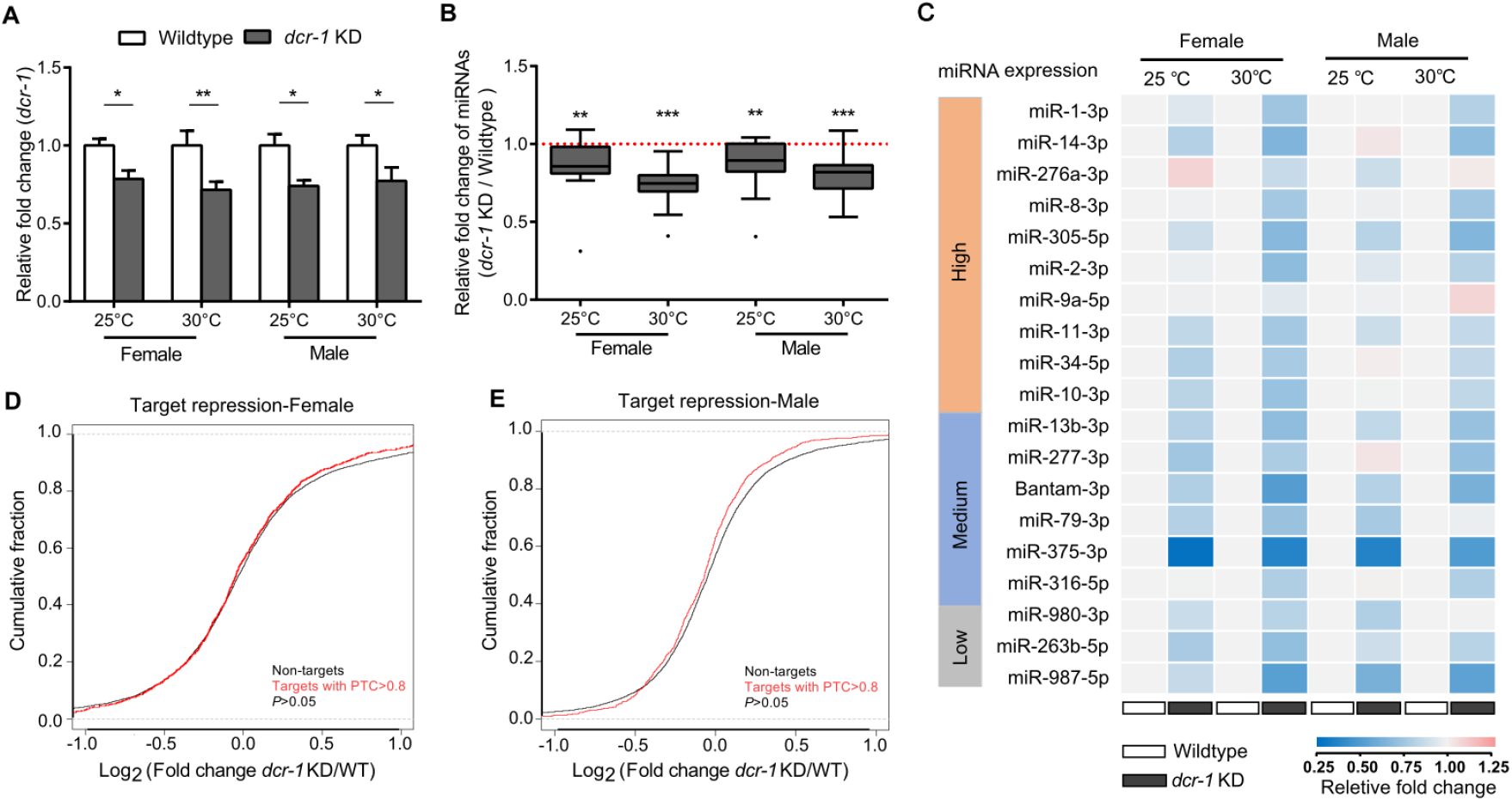
*dcr-1* KD leads weak but broadly reduction in miRNA pool. A) Impact of RNAi on *dcr-1* expression. Bar plots show the mean with SD for the relative expression of *dcr-1* in female and male L3 with different genotypes at two temperatures. For both sex and temperature, *dcr-1* expression significantly decreases 25-30% in *dcr-1* KD larvae. (Two-tail Student’s *t* test, \**p*<0.05, \*\**p*<0.01.) B-C) Impact of *dcr-1* RNAi on miRNA expression. (B) Summary of relative fold change of 19 selected miRNAs, accounting for over 50% of the total sequencing reads in the L3 (for more details, see **Fig.S1**). These miRNAs are down-regulated modestly but broadly at both 25°C and 30°C, one sample *t* tests are performed to test whether the up-regulation is significantly deviates from 1, ** *p*<0.01, *** *p*<0.001. (C) Heat map shows relative expression of selected miRNAs in female and male L3 with different genotypes at two temperatures. Relative expression levels of miRNAs in wildtype flies are set to 1. D-E) Impact of *dcr-1* RNAi on miRNA targets in female (D) and male (E) transcriptomes. Conserved miRNA targets (TargetScan PTC>0.8) do not show significant change compared to non-targets. (*p*>0.05, Kolmogorov–Smirnov test). QRT-PCR is performed in triplicate for genes and miRNAs, *rp49* and *U6* are used as inner reference controls for genes and miRNAs, respectively. Genotype: Wildtype (*T98-Gal4/+*;*UAS-GFP/+*); *dcr-1* KD (*T98-Gal4/+*;*UAS-Dicer-1* ^*HMS00141*^ */+*).

As a consequence of the Dicer-1 knock-down, most miRNAs are slightly down regulated, usually in the range of 15% - 25% (**Fig. 1B** and **Fig. S1**). **Fig. 1C** presents the expression differences of individual miRNAs in the *dcr-1* KD background. These selected miRNAs range from being highly to lowly expressed and the level of reduction appears even across the board. Because the complete knockout of each miRNA would generally increase the target gene expression by < 50%, the partial knockdown is expected to have rather mild effects on these expression levels **Fig. 1D** and **1E** corroborate the expectation. **Fig. 1** panels collectively show that *dcr-1* KD exerts weak but broad perturbations on the transcriptome.

### PART II-Developmental effects of *dcr-1* KD

#### Phenotypic consequence of dcr-1 KD under standard culture conditions

During the development, GRNs are constantly perturbed by external and internal noises. Although GRNs must have the capacity to attenuate the perturbations, the consequence is expected to be cumulative, affecting the late stages of development most severely. We therefore measured the life-history traits (**Fig.2A**) across the developmental stages (egg hatchability, larval viability, pupal viability and adult longevity) expecting normal development in the early stages but uncertainties in late ones.

**Figure.2.**
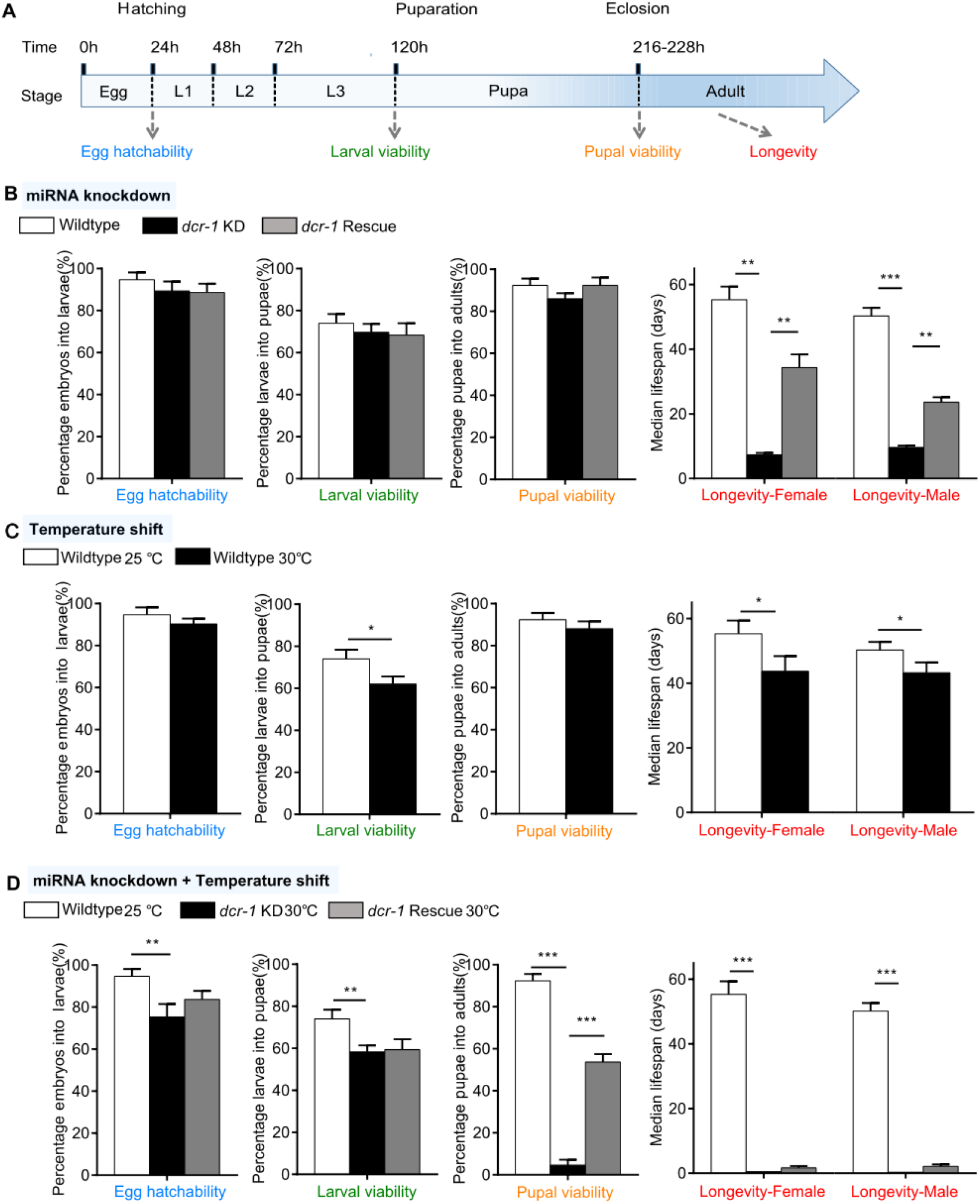
Phenotypic consequences of *dcr-1* KD under standard and high temperature. A) Experimental design for fitness component assays (indicated by arrows) across the whole *Drosophila* developmental process. Four typical fitness components (egg hatchability, larval viability, pupal viability and adult longevity are measured. B) Phenotypic consequences of miRNA knockdown. Compared with wildtype, *dcr-1* KD files reveal no significant changes in egg hatchability, larval or pupal viability while strikingly decrease longevity at 25°C. This phenotypic defect can be partially rescued by co-expressing *UAS*-*dcr-1* in the rescue line. (One way ANOVA with Tukey correction for multiple comparisons, ** *p*<0.01, *** *p*<0.001). C) Phenotypic consequences of temperature shift. High temperature slightly reduces these fitness components. Student’s *t* test is used for significance tests, * *p*<0.05. D) Phenotypic consequences of both temperature shift and miRNA knockdown. *dcr-1* KD flies reveal striking decreases in pupal viability and longevity at 30°C. These phenotypic defects can be partially rescued in the rescue flies. (One way ANOVA with Tukey correction for multiple comparisons, * *p*<0.05, ** *p*<0.01, *** *p*<0.001). Genotype: Wildtype (*T98-Gal4/+*; *UAS-GFP/+*); *dcr-1* KD (*T98-Gal4/+*; *UAS-Dicer-1* ^*HMS00141*^ */+*); *dcr-1* rescue (*T98-Gal4/+*; *UAS-Dicer-1* ^*HMS00141*^*/UAS-Dicer-1*).

Under standard conditions at 25 °C, *dcr-1* KD flies indeed develop normally showing little adverse effects in egg hatchability, larvae viability and pupation. Strikingly, toward the very end of the development, adults experience 80% reduction in longevity in both sexes (One way ANOVA, p<0.05; **Fig.2B**). The phenotypic defect in lifespan could be rescued by reintroducing UAS-*dcr-1* into the *dcr-1* KD background (Methods and Materials, **Fig. S2**). The occurrence in the *dcr-1* KD flies of defects only in the late developmental stages is consistent with the hypothesis of miRNAs role in developmental canalization.

#### Phenotypic consequences of dcr-1 KD under high temperature

We then explored if miRNA can ameliorate the perturbation of temperature shift from the standard 25 °C to 30 °C. The higher temperature, known to be stressful to the flies(Chen et al. 2015; Sgro et al. 2016; Vihervaara et al. 2018), should not be uncommon in nature. In the *dcr-1* KD background, the perturbation happens in flies with the stability control compromised, analogous to a high-wire walker without the pole.

Growing in 30 °C, *dcr-1* KD flies displayed severe phenotypic defects with more than 95% decrease in both pupal viability and adult longevity (**Fig. 2C vs 2D**). Hence, the defects are much more severe than in 25°C. Most flies cannot complete the normal development which could be stalled in various pupal stages (**Fig S3**). Since the failure to develop does not happen in a defined stage, the observation again suggest some cumulative effects in the miRNA KD background. In short, even though fly development is well canalized, the modest miRNA knockdown still leads compromised stabilization (**Fig. 2D**).In all experiments, the rescue line is also used. Although the rescue can usually be observed, it may at times be rather weak under extreme conditions as in **Fig. 2D**. We will discuss this incomplete rescue in Discussion.

#### Phenotypic consequence of dcr-1 KD under transient high-temperature shift

The cumulative effect of miRNA KD on development can be best demonstrated with transient temperature shift. The temperature is raised to 30°C in either a 24- or 72-hour pulse (**Fig. 3A**). We found *dcr-1* KD could significantly decrease fly fitness by all heat pulses (**Fig. 3B – 3C**; One way ANOVA, overall *p*<0.05).

**Figure.3.**
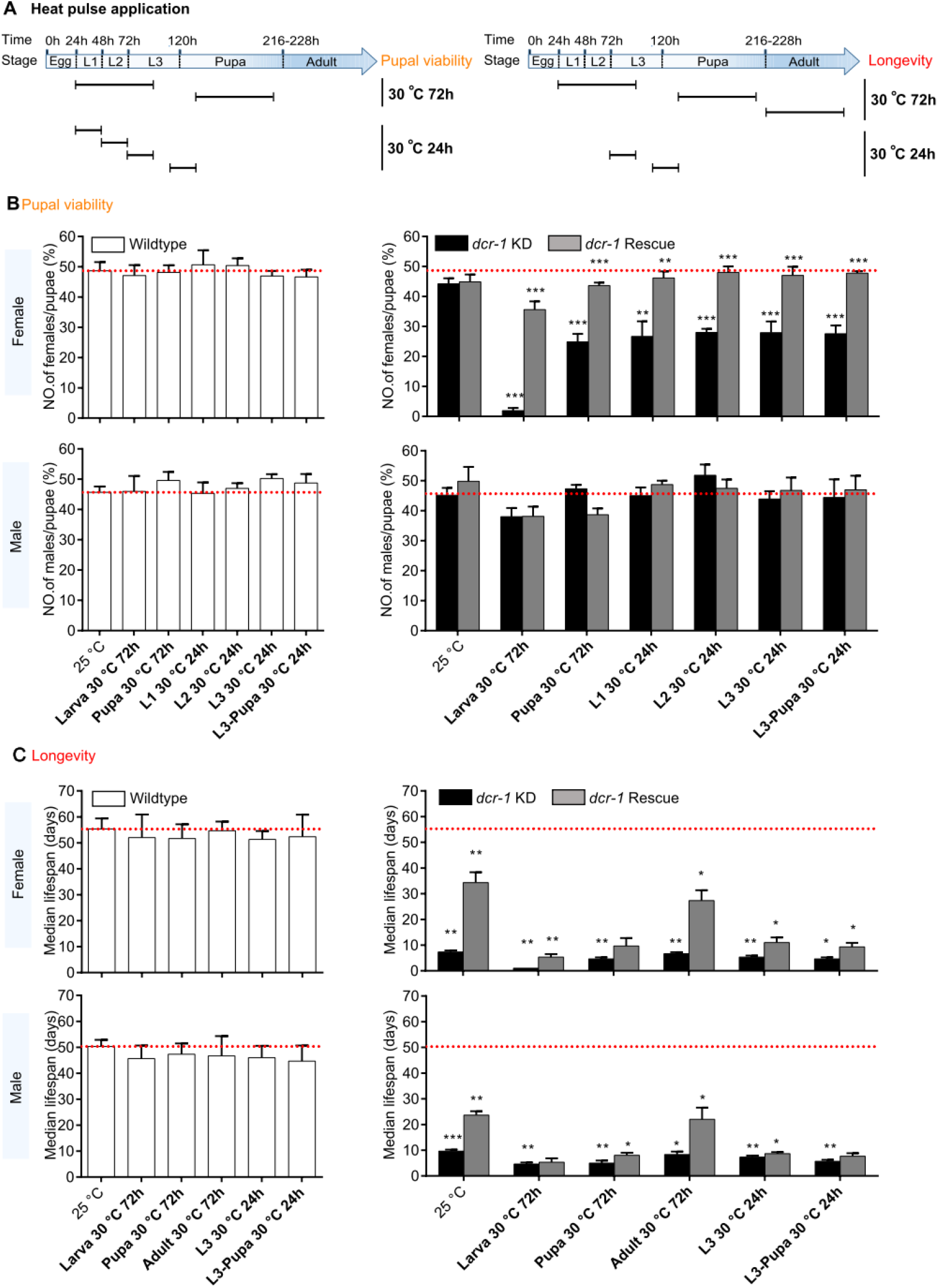
Phenotypic consequences of *dcr-1* KD after transient temperature shifts. A) Experimental design of the transient temperature shift (heat pulse) assay. Segment lengths indicate heat pulse durations (72h or 24h). Two typical fitness components, pupal viability and longevity, are measured. B) Pupal viability of wildtype, *dcr-1* KD and *dcr-1* rescue flies after different transient temperature shifts. Left panel, pupal viability of wildtype flies under different heat pulse assays. Right panel, pupal viability of *dcr-1* KD and *dcr-1* rescue flies under different heat pulse assays. The red line indicates the wildtype value without perturbation. After different heat pulse assays, pupal viability of wildtype flies decreased slightly, while pupal viability of *dcr-1* KD flies decreased significantly, especially in females. The phenotypic defect overall can be partially rescued in the rescue flies. Note that larval stages of *dcr-1* KD female flies are more sensitive to heat pulses (reduce to ∼1.9% while others were around 26% compared with wildtype) and show cumulative effect (27% (L1 24h HP) × 28% (L2 24h HP) × 28% (L3 24h HP) = 2.1%, which is nearly the same proportion, 1.9% with larva 72h HP) of the lethality. (One way ANOVA with Tukey correct for multiple comparisons is performed, ** *p*<0.01, *** *p*<0.001). C) Longevity of wildtype, *dcr-1* KD and *dcr-1* rescue flies. After different heat pulse assays, lifespan of wildtype flies decreased slightly, while lifespan of *dcr-1* KD flies decreased significantly. The phenotypic defect can be partially rescued in the rescue flies. For *dcr-1* KD flies, 72h heat pulse at the larval stage has the strongest effect in both sexes. Note that *dcr-1* KD females are more sensitive to heat pulses than males (One way ANOVA with Tukey correction for multiple comparisons, * *p*<0.05, ** *p*<0.01, *** *p*<0.001).

For females, *dcr-1* KD flies reflect significant decreases in both pupal viability and lifespan after transient temperature shift. We observed that as the perturbations are shifted earlier (larvae vs pupae or adults), the affected stages are shifted correspondingly, leading to more severe decrease in both viability and longevity (**Fig. 3B&C**, up panel). We also found that these fitness decreases reveal cumulative effect as a function of developmental progression (**Fig. 3B&C**, up panel), with stronger effects toward the late developmental stages (adults). For males, *dcr-1* KD flies also show significantly decreases in lifespan after transient temperature shift (**Fig. 3B&C**, down panel). As for the sex differences in the phenotypes of *dcr-1* KD, we suggest that the differences may simply be a “Tail end” effect (a small difference in mean becoming a large difference in the tail of the distribution) reflecting the innate sexual difference. Collectively, these results suggest that the pronounced phenotypes of *dcr-1* KD at 30°C represent the cumulative effects of perturbations through development.

### PART III – Changes in the GRN of *dcr-1* KD

The phenotypic perturbations are likely the manifestations of changes in the underlying transcriptomes and GRNs. Following this logic, we hypothesize that, in the developmental stages when the phenotype is still normal but is at the cusp of imminent changes (like a faltering high-wire walker), the underlying transcriptome should be showing some deviations from the norm. We hence examine the transcriptome right before phenotypic abnormalities become observable, i.e., the late third instar larvae (L3).

The transcriptome analysis of **Fig. 4A** shows that the transcriptome in *dcr-1* KD appears normal in the absence of perturbation, thus echoing the phenotypic analysis. In **Fig. 4A**, correlation coefficients are organized in 3 tiers – wildtype vs. *dcr-1* KD (orange), 25 °C - vs. 30 °C (green) and male vs females (blue). In the comparisons, we observed sex to have the largest impact (coefficient ranging from 0.841 to 0.879) while the temperature effect is relatively modest (coefficient ranging from 0.976 to 0.984 and 0.938 to 0.978 for male and female, respectively). Most interesting, *dcr-1* KD has very little effect (between 0.975 and 0.988) on the transcription patterns. Further statistical analysis by a general linear model (Methods and Materials) reveals 56%, 32% and 9% of the genes are significantly affected by sex, temperature and *dcr-1* KD respectively (**Fig. S4**). When dividing sex and adding the interactions into the general linear model (Methods and Materials), we observed the consistent pattern that *dcr-1* KD has a smaller effect on the transcriptome compared to temperature shift (**Fig. 4B-C**). These results support the conjecture of a normal transcriptome in *dcr-1* KD in the absence of perturbation.

**Figure 4.**
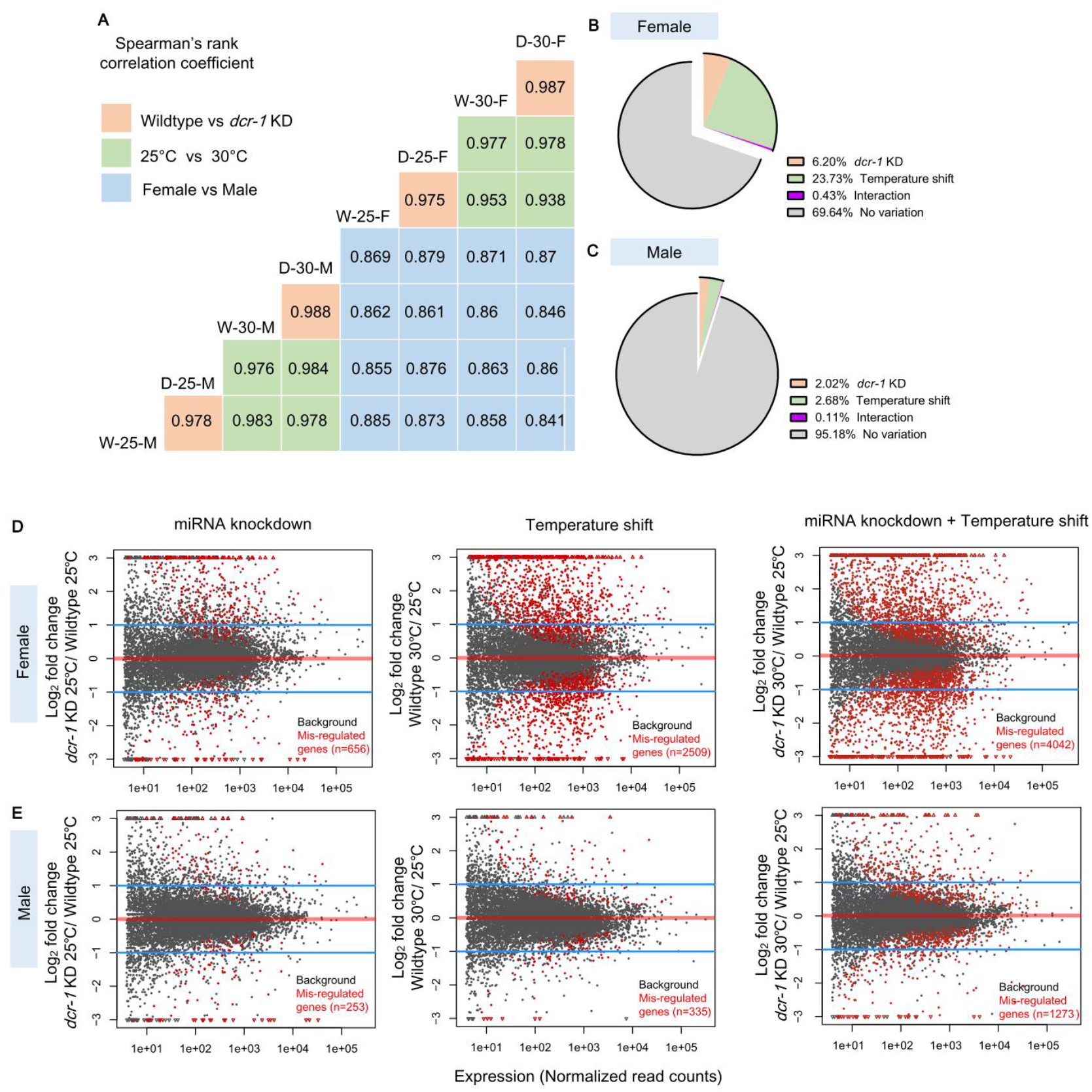
*dcr-1* KD results in “normal” but wobbly transcriptome. A) Pairwise correlations among whole transcriptomes in RNA-seq data sets. RNA-seq data was generated for each sex (female and male) ×genotype (Wildtype and *dcr-1* KD) ×temperature (25°C and 30°C) combination for L3.Correlations fall into three tiers, reflecting the impact of sex (highlighted by blue), temperature (highlighted by green), and *dcr-1*KD (highlighted by orange), respectively. B-C) Impact of temperature, *dcr-1* KD and their interaction on gene expression profiles for females (B) and males (C).The pie plot shows the percentage of genes affected by *dcr-1*KD, temperature shifts, genotype ×environment interactions and no variation according to a generalized lineal model method (see methods). D-E) Significantly dysregulated genes affected by three types of perturbation for females (D) and males (E). Genes affected by *dcr-1* KD, temperature shift and both are plotted in left, middle and right panel, respectively. Grey dots show all genes expressed in L3, while red dots show the significantly dysregulated genes detected by DEseq2 with FDR<0.1.

To see if the modest reduction of miRNA pool will reduce GRN stability, we compared the number of dysregulated genes under three different comparisons: *dcr-1* KD, temperature shift and both. We observed that despite *dcr-1* KD has much less impact on transcriptome, *dcr-1* KD exacerbates the stress of temperature shift on the transcriptome, resulting in more dysregulated genes **(Fig.4D-E)** in various gene ontologies **(Table S1)**. In conclusion, GRN becomes less stable with more dysregulated genes when miRNAs are modestly reduced across the board.

## Discussion

In 1942, Waddington proposed that “developmental reactions … bring about one definite end-result regardless of minor variations” (Waddington 1942). This statement is the genesis of the canalization view. Since then, the genetic circuitry required for canalization has been a constant topic. The view on canalization in recent years has been extended from the analysis of small-scale circuits (Ebert and Sharp 2012; Siciliano et al. 2013), to modules (Peláez and Carthew 2012), and then to large-scale regulatory networks (Ambros 2019; Chen et al. 2019). In particular, Chen et al. (2019) adopted the May-Wigner theory (May 1972; May 1974) on the stability of complex systems to show how miRNA actions can stabilize GRNs.

Obviously, developmental canalization requires GRN stability such that an unstable GRN would likely bring about un-canalized development. However, the connection between GRN stability and miRNA actions does not necessarily connect developmental canalization with miRNAs. An attempt at this connection has been reported by Liufu et al. (2017) that the same miRNA may regulate the same phenotype via multiple target genes, sometimes in the same but often in opposite directions. In other words, the miRNA-target gene relationships are highly redundant and incoherent.

In this study, the connection between miRNA actions, GRN stability and developmental canalization is established. We should note that, in previous reports (Li et al. 2009; Cassidy et al. 2013; Kasper et al. 2017) including the study by Liufu et al. (2017), the evidence is based on a few chosen miRNAs. This study provides the first empirical evidence supporting the theoretical prediction that miRNAs collectively serve as the canalization agent of development. Given that the environment fluctuates constantly, but often mildly, our results show how development may cope with such most common perturbations.

## Materials and methods

### Fly stocks

The *UAS-GFP* line (w1118/Y; *miniwhite-UAS-eGFP*/*miniwhite-UAS-eGFP*) was generated by our lab. It is a red-eyed strain with the insertion of *mini white* and *UAS-eGFP* at 2 chromosome 51D using the PhiC31 site-specific chromosomal integration system. T98-Gal4, *UAS-Dicer-1* ^*HMS00141*^, and *UAS-Dicer-1* were obtained from the Bloomington stock center and are described in flybase (http://flystocks.bio.indiana.edu/), stock numbers are 6996, 34826 and 36510, respectively. All flies were raised at 25°C on a standard sugar-yeast-agar medium and under 12:12 hr light/dark cycles. Heat treatment was carried out in the incubator at 30°C for specified times.

### Fly viability assay

Embryos were collected from 6-9 day-old flies, over 1-2h on grape juice agar plates. Before embryo collection flies were kept in bottles with frequently replaced grape juice agar plates for 3 days to eliminate old eggs.

For fly viability presented in **Fig.2**, three batches of 100 to 200 embryos were collected for each genotype and treatment (three genotypes ×two treatments, *dcr-1* KD, wildtype and *dcr-1* rescue; 25°C, 30°C). At day 0, the number of first instar larvae (L1) was counted after 30 h, the total number of pupae and adults was counted at day 11-12. This schedule was used for flies raised at both 25°C and 30°C. Egg hatchability (embryo viability) = L1 number/ egg number; larval viability = pupa number / L1 number; pupal viability (eclosion rate) = adult number/ pupa number.

For pupal viability presented in **Fig.3**, three batches of 100 to 200 embryos were collected for each genotype and treatment (three genotypes × seven treatments, *dcr-1* KD, wildtype and *dcr-1* rescue; 25°C and heat pulse shown in **Fig.3A** left panel). At day 0, the total number of pupae and adults (distinguishing females and males) was counted at day 11-12. Eclosion rate for female = adult female number/ pupa number; Eclosion rate for male = adult male number/ pupa number.

### Determining pupal lethality stages

To define pupal lethality stages (**Fig.S3**), we collected un-eclosed pupae at day 12 for each genotype and treatment (three genotypes × two treatments: *dcr-1* KD, wildtype and *dcr-1* rescue; 25°C and 30°C). Metamorphosis of these pupae stopped at specific stages. 100 to 200 wildtype flies were counted for each phenotype. Pupal development stages are according to (Bainbridge and Bownes 1981). Briefly, pupae with yellow eyes are regarded as dead at late pupalstages, the rest at early stages; for the early pupae, pupae with legs and wings achieving full extension along the abdomen are regarded as dead at P5 to P7, the rest at P1to P4. Late pupae with mature bristles on most abdomen regions are regarded as dead at P14 to P15, the rest at P8 to P13.

### Adult longevity assay

Adult longevity (**Fig. 2&3**) assays were performed as described in (Chen et al. 2014). Briefly, virgin females and un-mated males were collected separately, three vials (replicates) of 20 flies of each genotype and treatment (three genotypes × seven treatments×two sexes: *dcr-1* KD, wildtype and *dcr-1* rescue; 25°C,3 0°C and heat pulse shown in **Fig.3A** right panel; female and male). Dead flies were counted every two to three days and flipped to fresh vials until all had died. Median survival was then determined and analyzed.

### Detecting miRNA and gene expression by RT-PCR

For each genotype and treatment (three genotypes × two treatments: *dcr-1* KD, wildtype and *dcr-1* rescue; 25°C and30°C), three batches of 6-10 female third instar larvae (L3) were collected independently as biological replicates. L3 sex was distinguished by gonad morphology. For each replicate, total RNA was extracted using the Ambion TRIzol^®^ Reagent (code No. 15596018). Total RNA was reverse-transcribed into cDNA using the TARAKA Mir-X™ miRNA First Strand Synthesis Kit (code No.638315) and the TOYOBO ReverTra Ace qPCR RT Master Mix with gDNA Remover (Code No. FSQ-301) for miRNA and protein coding gene analyses, respectively. RT-PCT was performed on an ABI PRISM 7900 sequence detection system, TARAKA SYBR^®^ Premix Ex Taq™ (Tli RNaseH Plus) (code No.RR420) was used to detect amplification products. Relative expression levels were calculated as 2 ^−ddct^. Primers used in this study are shown in TableS1. *rp49* and *U6* were used as internal reference control for protein coding genes and miRNAs, respectively.

### Small RNA-seq analyses and target site prediction

Small RNA libraries of L3 were retrieved from GEO (Gene Expression Omnibus) with accessions GSM322208 and GSM322245. High confidence miRNA precursors and mature sequences were retrieved from miRBase Release 21 (http://www.mirbase.org) (Kozomara and Griffiths-Jones 2014). *Drosophila melanogaster* genome and 3’UTR sequences (r6.04) were retrieved from FlyBase (http://flybase.org/) (Attrill et al. 2016). The expression of each mature miRNA was measured using miRDeep2 version 2.0.0.7 (Friedlander et al. 2012) with default parameters, normalized by total reads matching all miRNA precursors per library and scaled as Reads Per Million (RPM). miRNA targets were predicted using TargetScan Fly version 7.2 (Agarwal et al. 2018). Conserved miRNA targets were predicted using TargetScan based on conservation (TSs) with PCT > 0.8 as previously described (Lu and Clark 2012).

### RNA-seq analyses

Total RNA was extracted from ∼10 larvae using the TRIzol^®^ Reagent, ribo-depleted total-RNA libraries were constructed and sequenced on Illumina HiSeq 2000 at BGI (http://www.genomics.cn/index). *Drosophila melanogaster* genome (BDGP6.83) was retrieved from the Ensembl database. RNA-seq reads were mapped to the *D. melanogaster* reference genome using Tophat (v2.1.0). The number of reads mapping to each protein-coding gene were quantified using HTseq (v0.6.0). Significantly disregulated genes were called using DEseq2 (Love et al. 2014) with *FDR* < 0.1. Similar to previous work (Yeh et al. 2014), a Generalized Linear Model was used to calculate the impact of each factor (Sex, Temperature and Genotype) on gene expression using the DESeqDataSet object:

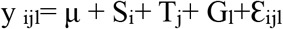

In this formula, y _ijl_ refers gene expression for the *i* th sex, *j* th temperature, *l* th genotype; μ is the baseline expression, S_i_ is the effect of the *i* th sex, T_j_ is the effect of the *j* th temperature, G_l_ is the effect of the *l* th genotype, ε is the error term. To define the genes affected by genotype (*dcr-1*KD), temperature, genotype × environment interactions for each sex, the following Generalized Linear Model was used:

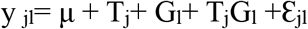

In this formula, y _jl_ refers gene expression for the *j* th temperature, *l* th genotype; μ is the baseline expression, T_j_ is the effect of the *j* th temperature, G_l_ is the effect of the *l* th genotype, T_j_G_l_ is the interaction term, ε is the error term.

Significantly dys-regulated genes affected by each factor were called with the *FDR* < 0.1 cutoff. To quantify gene expression and remove the dependence of variance on the mean, the function *vst* (variance stabilizing transformations) in DESeq2 was used to transform gene count data. Genes with read coverage less than 4 were not included in our analysis. GO enrichment was estimated using David 6.8 with Cutoff *P* value = 0.01 (Huang da et al. 2009). All raw sequencing data were submitted to the National Genomics Data Center (https://bigd.big.ac.cn/) with accession number PRJCA006274.

## Supplemental material

**Figure S1.**
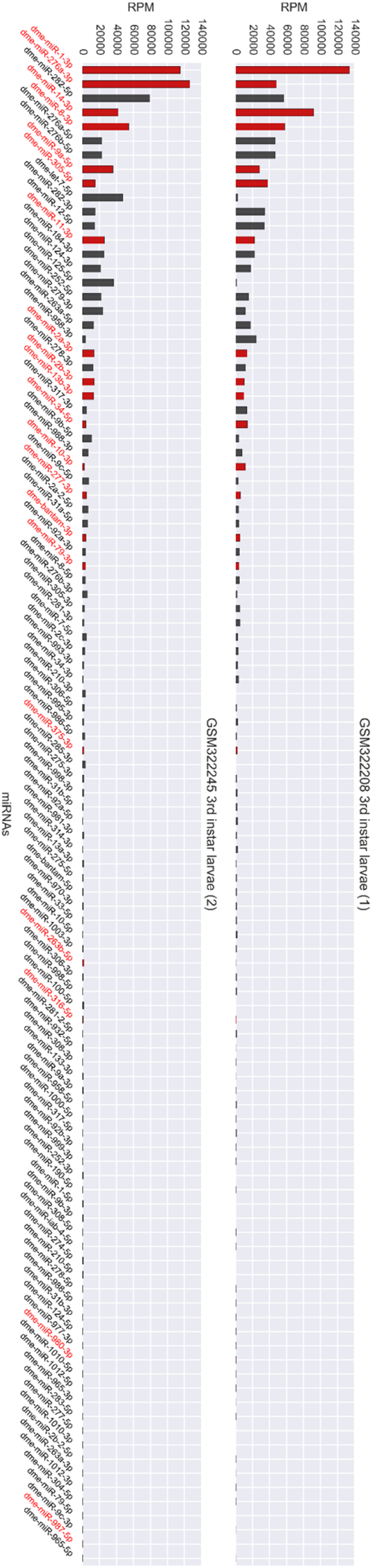
Normalized expression of all miRNAs (RPM) in L3. miRNAs surveyed in this study are highlighted in red.

**Figure S2.**
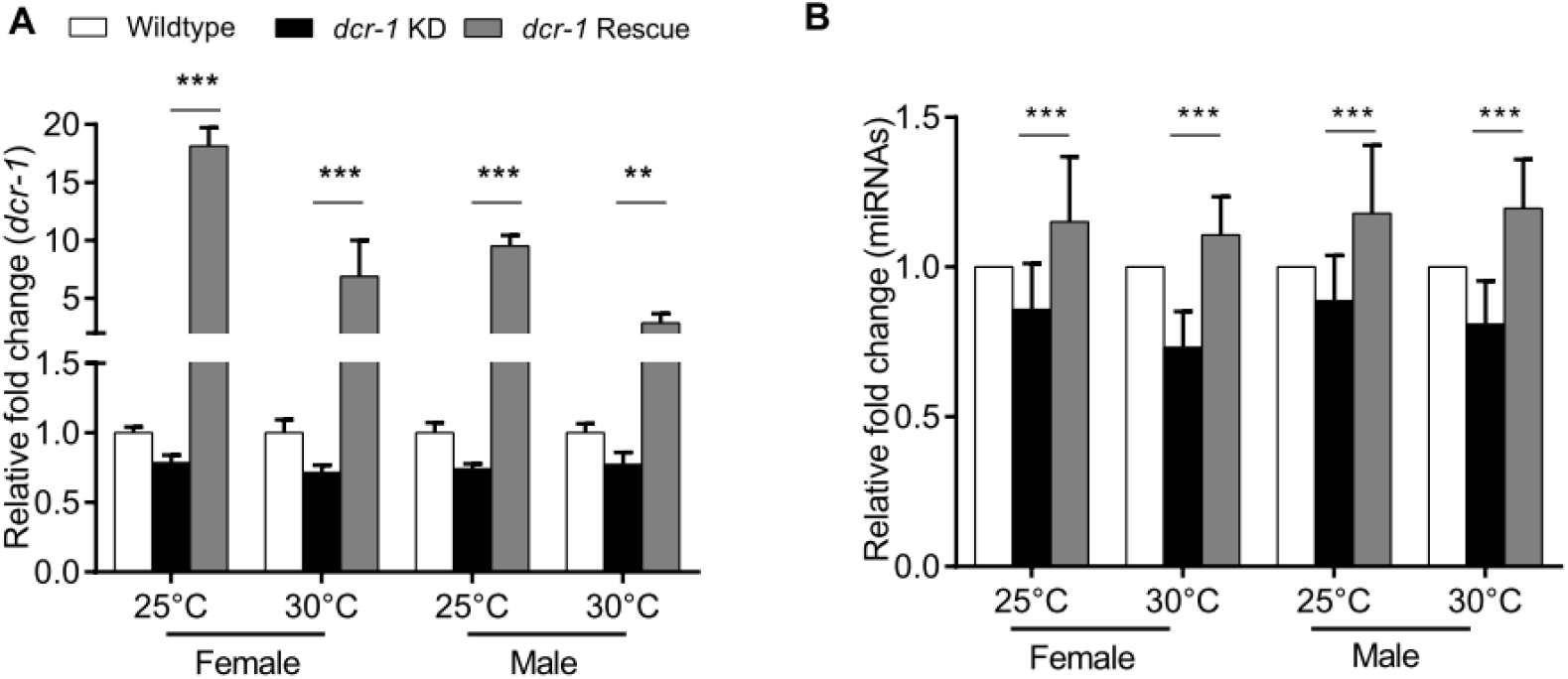
Impact of *dcr-1* rescue on the expression of *dcr-1* and the miRNA pool. A).Impact of *dcr-1* rescue on the expression of *dcr-1*. For both sex and temperature, down-regulation of *dcr-1* can be rescued by this construct. (Two-tailed Student’s *t* test, \*\**p*<0.01, \*\*\**p*<0.001) B) The impact of *dcr-1* rescue on the expression of miRNA pool. For both sex and temperature, the down-regulation of miRNA pool can be rescued by this construct. (Wilcoxon matched-pairs signed rank test, \*\*\**p*<0.001)

**Figure S3.**
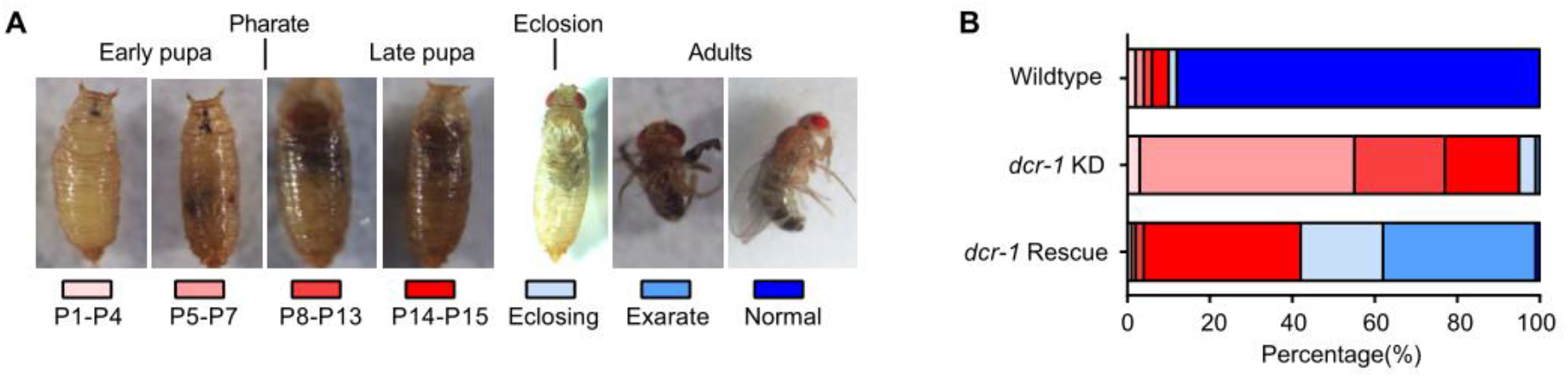
Pupal phenotypes of *dcr-1* KD at 30°C. A) *dcr-1* KD flies die at multiple pupal stages when raised at 30°C. B) Stacked bar plots show the percentage of flies at different stages. The lethality can be partially rescued by co-expressing UAS-*dcr-1*, especially for the early pupal stages (P5 to P7).

**Figure S4.**
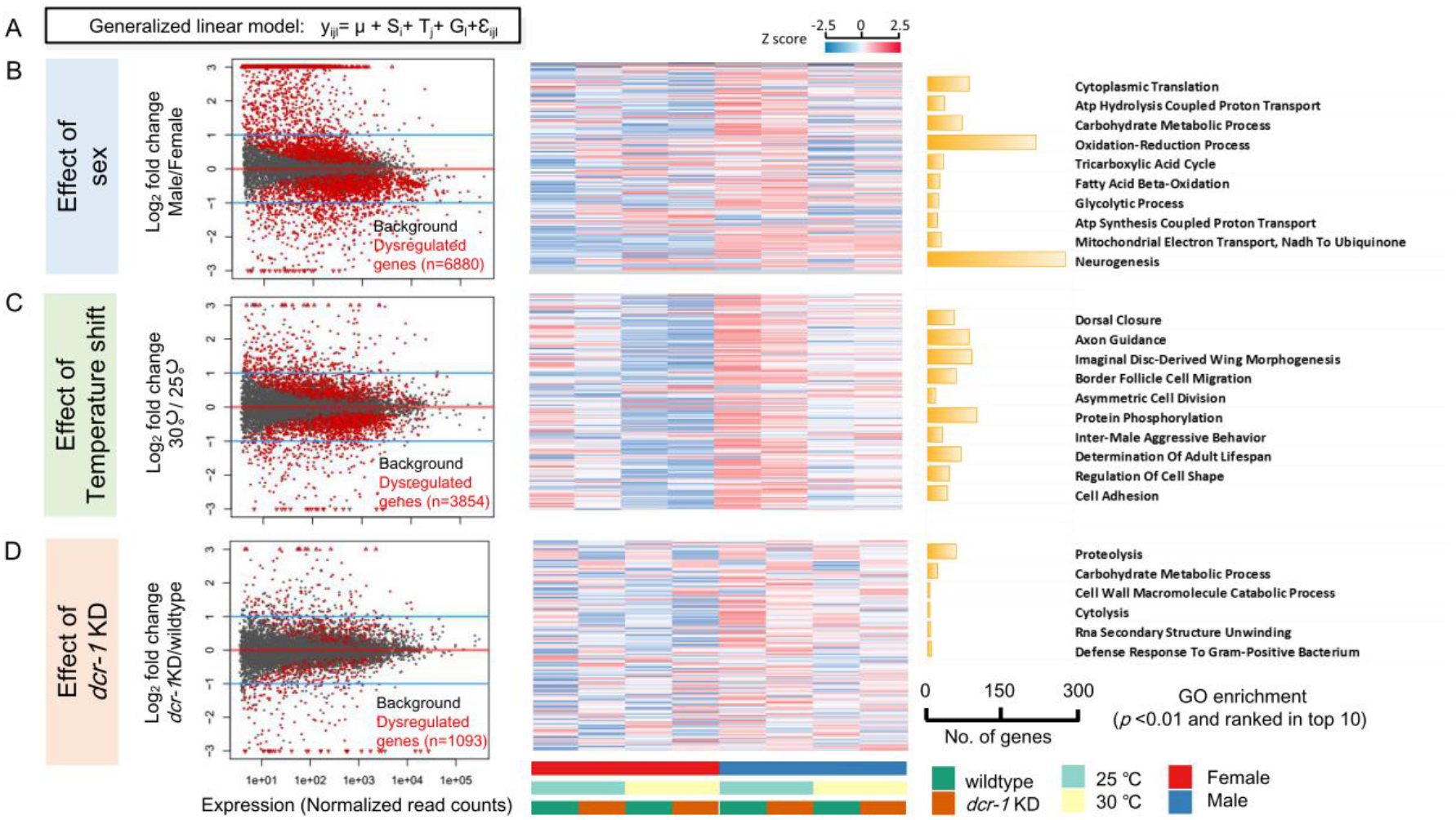
The effect of sex, temperature shift and *dcr-1* KD on transcriptional regulation. A) Calculation of the impact of each factor (Sex, Temperature and Genotype) on gene expression using a Generalized Linear Model (for details, see Materials and methods). B) Impact of sex on general transcriptional regulation. Left panel, genes significantly affected by sex across all data sets. Grey dots show all genes expressed in L3, red dots show the significantly affected genes detected by a generalized linear model at false discovery rate (*FDR*) <0.1. Middle panel, 150 genes affected by sex are randomly selected and presented by *Z* score. Right panel, GO enrichment, only the terms with modified Fisher Exact *p* value<0.01 and ranked in top 10 are shown. C) Impact of temperature shift on general transcriptional regulation. D) Impact of *dcr-1* KD on general transcriptional regulation.

**Table S1. GO enrichment for the genes affected by temperature shift, *dcr-1* KD and both.**

**Table S2.**
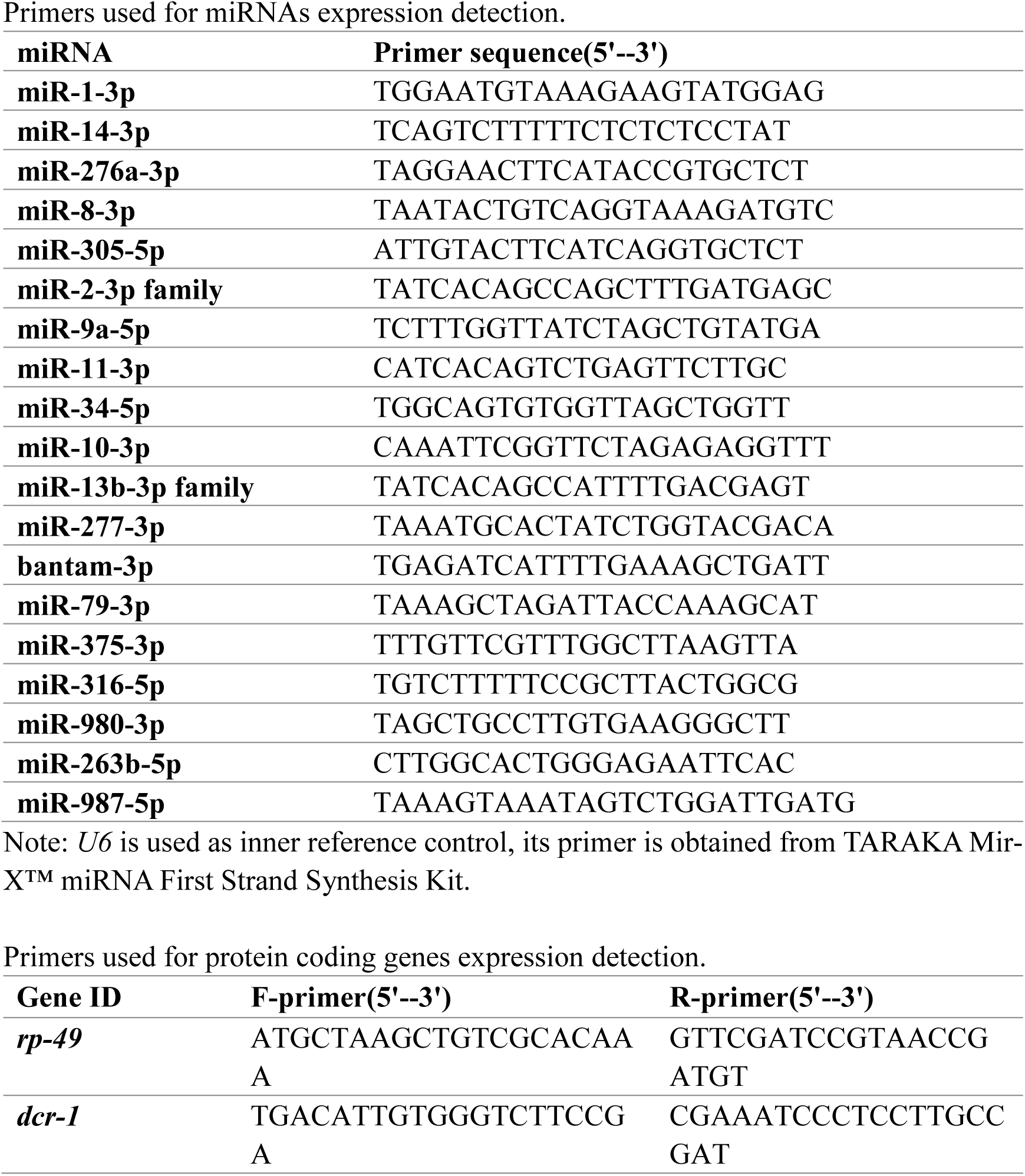
PCR primers used in this study.

## Declarations

### Ethics approval and consent to participate

Not applicable.

### Consent for publication

Not applicable.

### Availability of data and materials

All raw sequencing data were submitted to the National Genomics Data Center (https://bigd.big.ac.cn/) with accession number PRJCA006274. The open software used in this study is cited and materials can be requested from the authors.

### Competing interests

The authors declare that they have no competing interests.

### Funding

This work was supported by National Natural Science Foundation of China (31900417, 31730046, 31801081, 91731000), Guangdong Basic and Applied Basic Research Foundation (2019A1515010708, 2020A1515010467), and China Postdoctoral Science Foundation (2020T130748, 2020M672998).

### Authors’ contributions

G.A.L., Z.L., T.T., and C.-I.W. conceived and designed the study. G.A.L., Y.Z., and Q.C performed analyses with assistance from Z.T., and P.L. C.-I.W., G.A.L. and Y.Z wrote the paper, with critical revisions provided by all authors.

## Acknowledgements

We thank all members in the Wu laboratory for helpful comments and sharing of ideas. We thank Suhua Shi, Yongsen Ruan, Mei Hou, Yumei Huang and Haijun Wen for extensive discussions and critical reading of the manuscript.

## Notes

### Competing Interest Statement

The authors have declared no competing interest.

### Summary of Updates

Risive language errors.

